# Cell Rearrangement Generates Pattern Emergence as a Function of Temporal Morphogen Exposure

**DOI:** 10.1101/2021.02.05.429898

**Authors:** Timothy Fulton, Kay Spiess, Lewis Thomson, Yuxuan Wang, Bethan Clark, Seongwon Hwang, Brooks Paige, Berta Verd, Benjamin Steventon

**Affiliations:** Department of Genetics, University of Cambridge, Cambridge, UK; School of Biological and Behavioural Sciences, Queen Mary University of London, London, UK; The Alan Turing Institute, London, UK; Department of Zoology, University of Cambridge, Cambridge, UK; Centre for Artificial Intelligence, University College London, UK; Department of Zoology, University of Oxford, UK

## Abstract

As tissues elongate, cell rearrangement alters positional information in manner that must be accounted for to generate gene expression pattern. How this is achieved during paraxial mesoderm elongation is unknown. By reverse-engineering gene regulatory networks that predict single cell expression trajectories across the tissue, we find a network capable of recapitulating the full range of dynamic differentiation profiles observed both *in vivo* and *in vitro*. Simulating gene expression profiles on *in toto* cell tracking data sets reveal that temporal exposure to Wnt and FGF is generated by cell movement. The absence of reversal in gene expression towards a more premature gene expression state predicts the generation of aberrant tbx6 expression in the posterior progenitor zone that we then confirm by quantitative single cell imaging. Taken together, these results demonstrate cell rearrangement tunes the dynamics of mesoderm progenitor differentiation to generate pattern emergence as a function of temporal Wnt and FGF exposure.

## Introduction

During the elongation of the vertebrate body axis, rates of self-renewal and cell differentiation must be precisely coordinated such that a continual supply of progenitor cells are maintained through embryo elongation ***Henrique et al. (2015)***; ***Wilson et al. (2009)***. This temporal coordination is especially important within the presomitic mesoderm (PSM) where cells are segmented into somites at the anterior end of the tissue in a clock-like process ***Gomez et al. (2008)***. While much has been studied regarding the temporal dynamics of somitogenesis, less is known about the mechanisms that coordinate cell fate specification in the tailbud as cells become progressively specified to a PSM cell state ***Martin (2016)***. In zebrafish embryos, this transition is marked by a switch between tbx16 and tbx6 expression ***Warga et al. (2013)***; ***Nikaido et al. (2002)***. This pattern of gene expression appears strikingly robust at the tissue level and aligns with a graded decrease in both Wnt and FGF pathway activity ***Bajard et al. (2014)***; ***Spiess et al. (2022)***. Both morphogens generate a wavefront of positional information that ultimately times the differentiation of PSM cells and helps determine the dynamics of somitogenesis together with a Notch dependent clock ***Palmeirim et al. (1997)***; ***Dubrulle et al. (2001)***; ***Aulehla and Pourquié (2008)***; ***Aulehla et al. (2003)***.

While cells are undergoing coordinated differentiation, they are also undergoing a rapid series of cell rearrangement that are required as a driver of PSM elongation. In chick embryos, it has been shown that random cell rearrangements downstream of FGF signalling result in a coordinated posterior expansion of the tissue along the anterior-posterior axis ***Bénazéraf et al. (2010)***; ***Xiong et al. (2020)***. A similar transition from a liquid-like state of high cell motility in the posterior to a more solid-like state in the anterior has been observed in zebrafish embryos ***Mongera et al. (2018)***; ***Serwane et al. (2017)*** and is linked to an increased amount of cell rearrangement in the posterior progenitor zone ***Lawton et al. (2013)***; ***Das et al. (2019)***. Analysis of cell rearrangements in 3D further show how cells rapidly exchange their neighbours as the tissue compacts in the dorsal-ventral and medial-lateral axes and elongates along the anterior-posterior axis ***Thomson et al. (2021)***. How these movements impact the dynamics of PSM differentiation at the single cell level, yet still enable the generation of stable patterns of T-box expression at the tissue level is currently unknown.

Recent work has highlighted the extent to which the dynamics of the zebrafish somitogenesis clock is intrinsic to each cell. Previous studies had shown that cells cultured *in vitro* are capable of eliciting transient oscillations of Notch pathway activity, as assayed with reporters for her1 expression ***Webb et al. (2016)***. However, it was unclear whether such oscillations might be an emergent property of small groups of cells enabling the production of signalling dependent coordination between cells. More recently, it has been demonstrated that her1 oscillations can still be observed within isolated cultures of cells and in the absence of any signals added to the medium ***Rohde et al. (2021)***. Importantly, these cells slow down their oscillations and upregulate mesp expression, a marker of somite polarity ***Rohde et al. (2021)***; ***Yasuhiko et al. (2006)***. These results demonstrate that PSM differentiation is to a large degree cell autonomous in zebrafish embryos. Furthermore, they open the question of how the dynamics of an intrinsic timer can be modulated to match the cell specific dynamics of differentiation that occurs *in vivo* given individual rates of movement from the tailbud.

To investigate the cell-intrinsic mechanisms that time cell differentiation in response to external signalling, a common approach is to reverse engineer a minimal gene regulatory network (GRN) that is sufficient in its description of the system to recapitulate spatial distributions of gene expression observed *in situ* ***Jaeger and Monk (2014)***. Such an approach has been successful in understanding the dynamics of morphogen interpretation in a range of systems including the *Drosophila* blastoderm and vertebrate spinal cord ***Crombach et al. (2016)***; ***Verd et al. (2014)***; ***Sagner and Briscoe (2017)***; ***Kicheva et al. (2014)***. However, inferring GRNs that can recapitulate the emergence of gene expression patterns in 3D, and in the context of rapid cell rearrangements is challenging. We have recently developed a fitting and modelling framework that achieves this in the context of the zebrafish PSM through the mapping of gene expression patterns onto *in toto* cell tracking datasets. This enabled the inference of Approximated Gene Expression Trajectories (AGETs) ***Spiess et al. (2022)*** which were used to reverse engineer candidate GRNs. Here, we aim to ascertain whether a single network can predict the full dynamics of PSM differentiation in both an *in vivo* and *in vitro* context.

To determine the range of timings for the movement of cells out of the tailbud, we first probed the limits of their differentiation dynamics by culturing tailbud progenitors *in vitro* and assaying the proportion of tbx16/6 expressing cells at subsequent timepoints. This analysis revealed that cells *in vitro* are remarkably coherent in their dynamics of differentiation. This is unlike the situation *in vivo* where cells undergo a range of differentiation dynamics as a function of their time spent in the posterior progenitor domain, and so are distributed across a broad range of temporal trajectories. This analysis revealed that unlike *in vivo*, where cells are distributed across a broad range of temporal trajectories, cells *in vitro* are remarkably coherent in their dynamics of differentiation. This is unlike the situation *in vivo* where cells undergo a range of differentiation dynamics as a function of their time spend in the posterior progenitor domain. As each temporal profile of differentiation is linked to a distinct profile of Wnt and FGF exposure, we reasoned that this non-autonomous regulation of a cell-intrinsic timer might be sufficient to describe the fully dynamic profiles of PSM differentiation both *in vitro* and *in vivo*. To test this, we asked whether a single set of GRN parameters as previously reverse engineered from AGETs can recapitulate all observed dynamics. Indeed, we find a GRN that can predict *in vitro* and *in vivo* differentiation dynamics. Strikingly, this simple three gene network can generate accurate emergent gene expression patterns when simulated on real tracking data, given measured inputs of FGF and Wnt signal gradients. Furthermore, it predicts the presence of aberrant tbx6 expression within the progenitor zone. Taken together, our model proposes that cell movements themselves act to regulate the dynamics of intrinsic cell differentiation dynamics, to generate the emergence of gene expression pattern as a function of morphogen exposure.

## Results

### A cell intrinsic timer drives differentiation of only a subset of tailbud progenitors

It has been previously reported that when cells are explanted from the PSM of the zebrafish embryo it can trigger an intrinsic timer to differentiate into Mesp positive somitic mesoderm ***Rohde et al. (2021)***. To assay the autonomy of T-box expression changes in isolation, we aimed to measure tbx16 and tbx6 expression changes in single cells explanted from the posterior-most region of the tailbud. Explants were first taken from embryos previously injected with mRNA encoding a nuclear-targeted kikume construct where the posterior 25% of the tailbud had been labelled. This confirmed that the posterior-most region alone was being taken for dissociation and subsequent cell culture (Figure 1A). Explants stained for tbx16 and tbx6 showed that most cells express high levels of tbx16 with only a few cells expressing tbx6 (Figure 1B).

**Figure 1.**
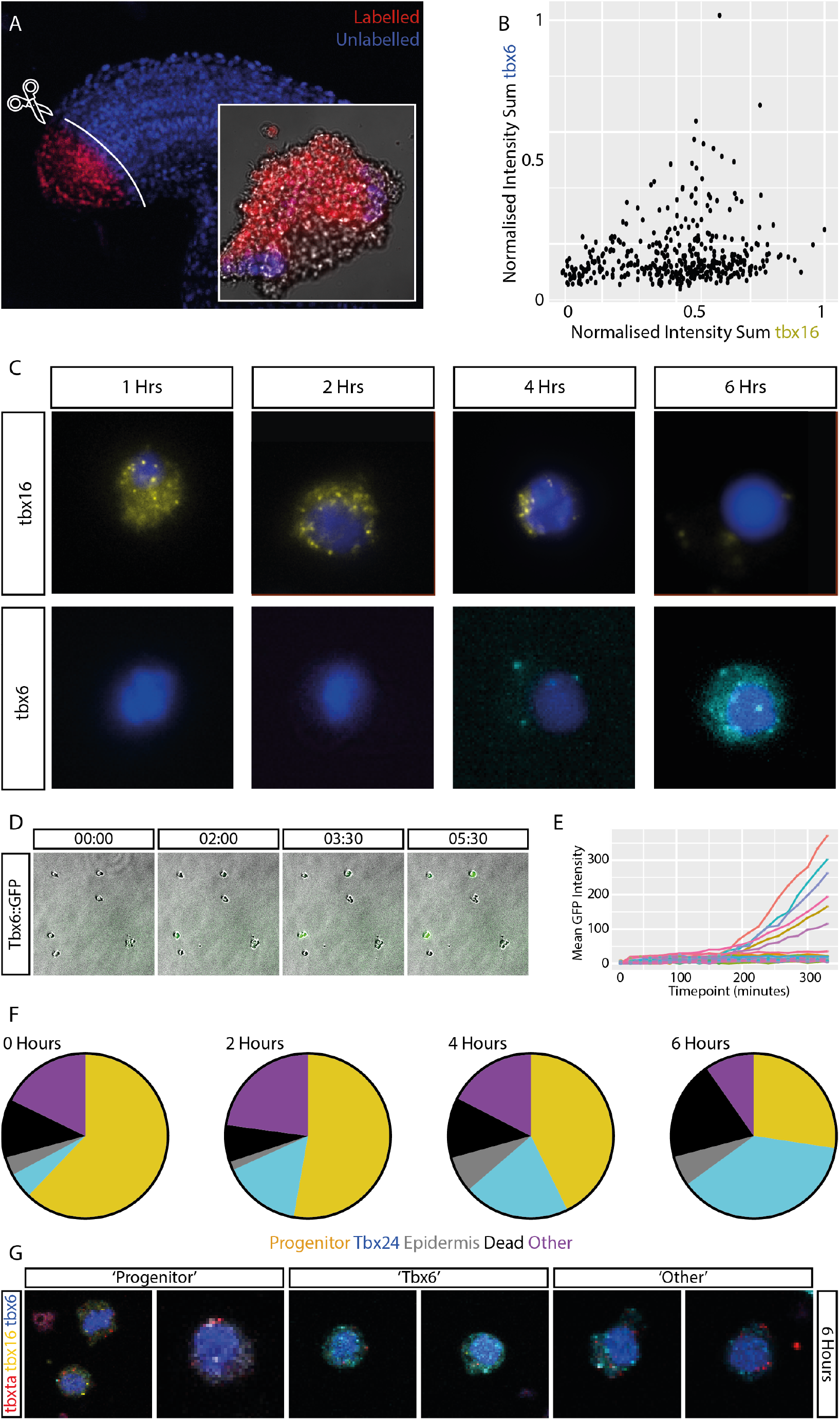
Presomitic Mesoderm progenitor cells differentiate *in vitro* forming a bimodal population of cells. **(A)** Labelling and dissection of the posterior 25% of the tailbud demonstrates explants are produced mostly of labelled cells. Explant shown in image insert. **(B)** From HCR data on whole explants, it is demonstrated that they are made of cells expressing tbx16 and only a small number of tbx6 expressing cells. By dissociation of these cells into single cells and culturing them in L15 media without supplementation. **(C)** cells are observed downregulating expression of tbx16 and increasing expression of tbx6 in fixed time points using HCR. **(D)** Using live imaging of a tbx6::GFP reporter, it was observed that GFP is observed only in a proportion of cells and that the **(E)** GFP signal appears synchronously in multiple cells after approximately 200 minutes *in vitro* culture. Cells not expressing GFP at this time do not express GFP even after 300 minutes of culture. Minimal to no cell divisions are observed occurring over the time course. **(F)** Through manual classification of single cell multiple HCR using tbxta, tbx16, tbx6, keratin18 probes, cells were defined as: “progenitor” (expressing tbxta and tbx16), “Tbx6” (expressing tbx6), “Epidermis” (expressing keratin18), “Dead” (with fragmented or unusual nuclei) or “Other” (none of the above gene expression patterns). It was observed the majority of cells begin the culture period as progenitors and that this proportion decreases as the proportion of tbx6 cells increases. The proportion of epidermis, other and dead cells stays relatively stable with some increase in dead cells by six hours. **G** Examples of Progenitor Cells, Tbx6 cells and cells classified as Other which express both tbxta and tbx6 together at the six hour time point.

Explants taken from the posterior tailbud were dissociated by gentle agitation in calcium and magnesium free PBS to produce single cells in suspension, and cultured in adherent culture under a media of defined L15 with no serum or signal factor supplementation, as previously described in ***Rohde et al. (2021)***. Under these culture conditions, cells downregulated expression of tbx16 and upregulated their expression of tbx6 over a period of 6 hours, as measured by HCR on fixed timepoint samples imaged at high magnification (Figure 1C). Further examination using a tbx6::GFP reporter ***Ban et al. (2019)*** revealed that GFP fluorescence increased in only a proportion of cells and that the GFP signal appeared synchronously after approximately 200 minutes (3.5 hours) in culture (Figure 1D,E).

Cells stained using multiplex HCRs for tbxta, tbx16, tbx6 and keratin18 at each time point were manually classified into five categories: progenitors, which expressed tbxta and/or tbx16, tbx6 positive cells expressing tbx6, epidermal cells expressing keratin18, dead cells which showed abnormal nuclei shape or fragmented nuclei, and other. Very few cells are observed dividing over this time course. Over time, the proportion of tbx6 positive cells increases at the expense of cells in the progenitor category (Figure 1F-G). These results together demonstrate that once isolated *in vitro*, cells from the posterior 25% of the tailbud have the potential to differentiate into tbx6 positive cells and that this differentiation occurs between 3 and 6 hours of culture. Furthermore, only a subset of posterior progenitors differentiate into tbx6 positive presomitic mesoderm, with the remaining cells staying in a tbxta positive progenitor state (Figure 1G).

### Cells *in vivo* differentiate over a range of time-scales as a function of their cell movement

We next aimed to relate the dynamics of T-box gene expression observed *in vitro* to the range of differentiation trajectories cells undertake *in vivo*. We first measured the time taken for a clone of 15-20 photo-labelled cells to spread from the from a tbxta positive progenitor region (Figure 2A) into a newly made somite (Figure 2C). We observed that in multiple embryos, the fastest cell entered a somite within 3 hours of being labelled in the posterior, with other cells exhibiting a range of longer times depending on how long they took to exit the progenitor domain.

**Figure 2.**
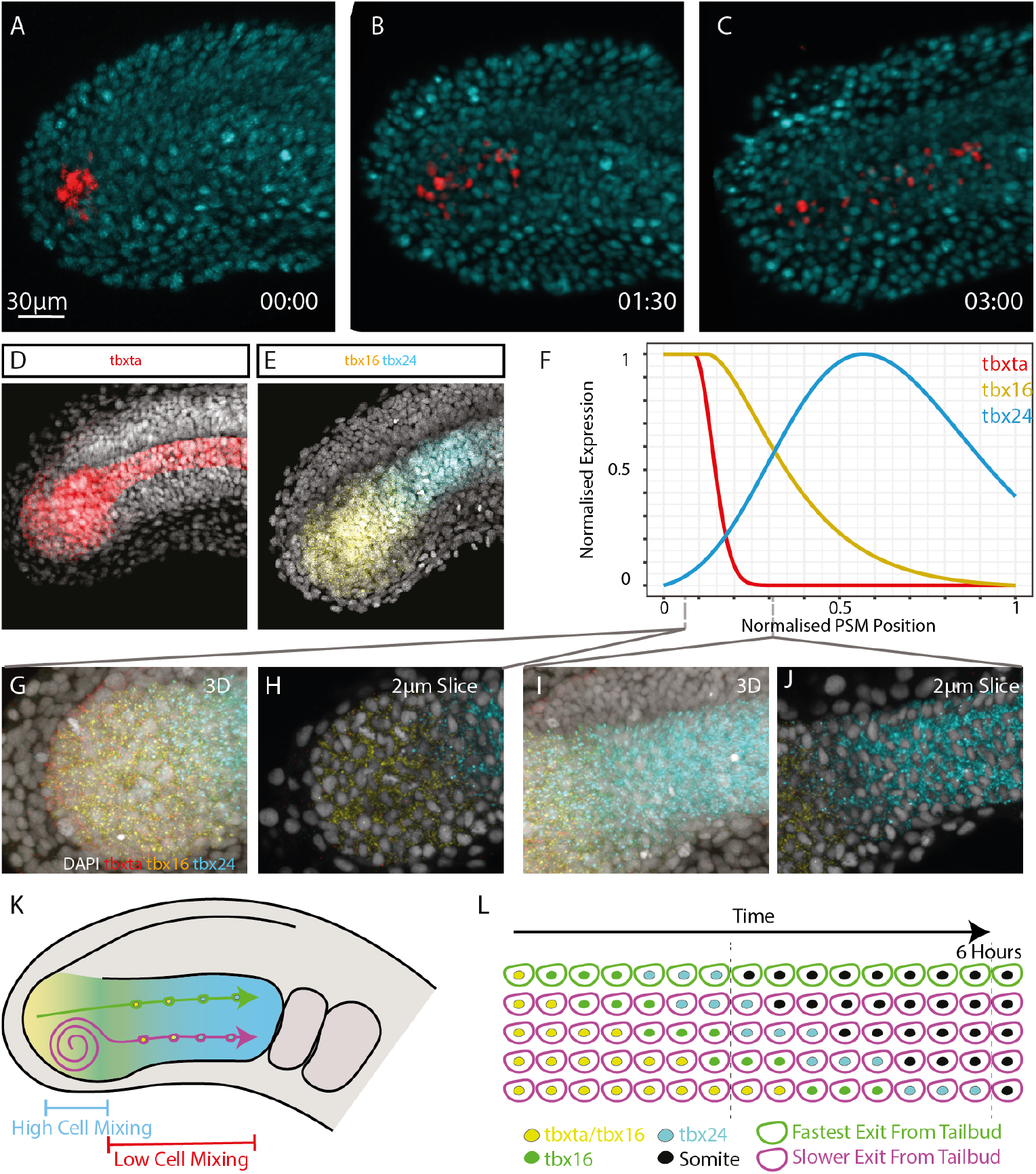
Stable patterns of T-box gene expression are formed across the presomitic mesoderm and tailbudd despite cell mixing. **(A-C)** Labeling of the posterior of the presomitic mesoderm demonstrates that cells spread throughout the entire tissue with the fastest cell entering the newly formed somite in 3 hours whereas other cells differentiate over a continuous range of timescales. **(D-E)** The T-box gene expression domains within the presomitic mesoderm are observed to be clear and stable using HCR against tbxta, tbx16 and tbx6. **(F)** These domains are quantified and plotted along a normalized posterior (0) to anterior (1) axis where 1 represents the posterior boundary of the most recently formed somite. Intensity of signal was also normalized between 0 and 1 from multiple embryos plotted together. Zoom in images of the **(G-H)** posterior tbx16/tbx6 transition domain and **(I-J)** anterior tbx6 domains as both 3D views and single 2µm slices. **(K-L)** Together this generates a situation where cells are continually mixing in the posterior presomitic mesoderm and differentiate over a range of timescales to become tbx6 positive, yet the domains of T-box gene expression remains stable.

To relate these cell movements to transitions in T-box gene expression, we used quantitative HCR to stain for tbtxa, tbx16 and tbx6 mRNA (Figure 2D-E). We masked the notochord and notochord progenitor cells and removed this tbxta signal from the image so that only the tbxta expressed within the PSM remained. The intensity of the HCR signal was then plotted across the PSM, normalizing the axis (where 0 represents the posterior end of the tailbud and 1, the posterior-most boundary of the most recently formed somite) (Figure 2F). This made it possible to quantify the expression patterns of the three T-box genes across the presomitic mesoderm and tailbud (Figure 2H-J). Our analysis revealed that cells undergo a range of temporal trajectories in gene expression, with the fastest cells transiting through to a newly formed somite in 3 hours; half the time taken for cells to fully upregulate tbx6 *in vitro* (Figure 2K-L).

### A reverse-engineered GRN recapitulates tissue-level T-box pattern formation and *in vitro* T-box gene expression dynamics as an emergent property of varying signalling dynamics

We have observed that tailbud progenitors can differentiate in a range of timescales in both *in vitro* and *in vivo* contexts. To investigate the underlying mechanisms that enable this, we set out to reverse-engineer a GRN that could be interrogated in manner that links intrinsic gene-regulatory dynamics with both cell movement and morphogen exposure. GRNs formulated as dynamical systems have been very helpful elucidating patterning mechanisms (see for example ***Balaskas et al. (2012)***; ***Raspopovic et al. (2014)***; ***Verd et al. (2018)*** among many others), however existing methodologies to infer GRNs from quantitative gene expression data have bypassed the role of cell movements in patterning processes (see Materials and Methods, and ***Spiess et al. (2022)*** for an in depth discussion). For this reason we set out to develop a methodology that would allow us to infer GRNs driving pattern formation in tissues undergoing morphogenesis which explicitly accommodated cell rearrangements and movements and applied it to the developing zebrafish PSM ***Spiess et al. (2022)***.

In brief, in order to retain cell movements we simulate GRNs directly on the cell tracks, obtained from *in toto* live imaging (Figure 3D), by formulating a GRN in each cell and simulating all of these in parallel using quantitative measurements of signal activity (Figure 3A-C) (we refer to this as live-modelling due to its parallel to live-imaging). This method allows us to observe the emergence and maintenance of pattern at the level of the tissue from the dynamics of gene expression being simulated in the single cells (Figure 3E-F), while still taking into account cell rearrangements explicitly. To reverse-engineer GRNs from our HCR data we constructed approximated gene expression trajectories (AGETs) by projecting the quantitative gene expression data onto each frame of the time lapse. We then used a small randomly spaced subset of 10 AGETs to infer candidate GRNs that would recapitulate T-box gene expression dynamics in single cells along the PSM. Candidate networks were then simulated on all the tracks allowing us to observe pattern formation on the developing PSM (Figure 3F) (see Materials and Methods, or refer to the original methodology paper ***Spiess et al. (2022)***).

**Figure 3.**
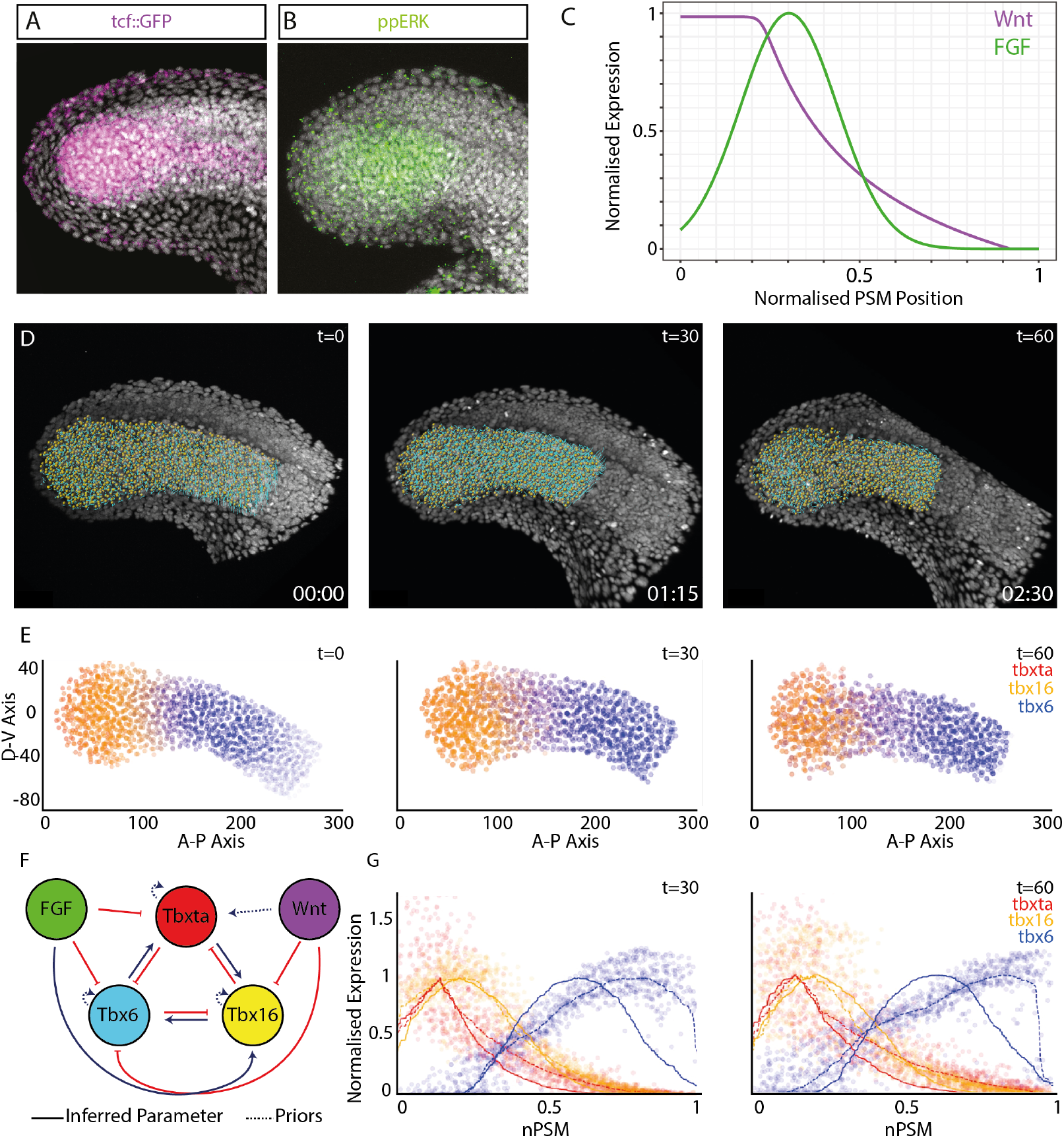
A Reverse Engineered Gene Regulatory Network Can Recapitulate *in vivo* Patterns of T-box Gene Expression With Cell Movements. **A-B** Measurements of the signalling environments within the PSM were made by either measuring the level of gfp mRNA produced from a TCF::GFP reporter to measure the level of Wnt signalling, or via an antibody stain against diphosphorylated ERK to measure FGF signalling. **C** These images were quantified along a normalised PSM axis as in Figure 2F. **D** We obtained live imaging data of the developing zebrafish PSM and tracked individual cells during 02:30hrs. t=0 shows the first frame in the time lapse where every nucleus in the PSM is highlighted by the tracking algorithm with a spot. t = 30 shows the 30th time frame which corresponds to 01:15hrs into the time-lapse and t = 60 shows the 60th frame in the movie, which corresponds to the end of the time lapse at 02:30hrs. Data from ***Thomson et al. (2021)***. In these tracking data, newly formed somites were deleted as soon as the morphological somite is formed. Non-presomitic mesoderm tracks were deleted from the entire movie at all time points. **(E)** The tracking data in (D) were used to implement the live-modelling framework where reverse-engineered zebrafish T-box GRNs are simulated on every cell track represented *in silico*. The live-modelling framework allows us to observe how tissue-level patterning emerges as a function of GRNs acting at the single cell level as the PSM undergoes morphogenesis. Tbxta in red, Tbx16 in yellow and Tbx6 in blue. **F** Reverse-engineered T-box GRN. **G** Quantified simulated gene expression patterns compared to the gene expression patterns measured in the embryo, previously quantified using HCR. Each dot represents the simulated concentration of a TBox gene in a single cell. Curves (dotted lines) were fit and normalised to the simulated gene expression and compared to the quantified experimental data (solid lines) at different time points (t=30 and t=60 shown here) to assess the goodness of fit.

Networks were inferred from fitting parameters to the data from 10 random single cell AGETs. We selected and clustered the networks that were able to recapitulate tissue-level T-box patterning when simulated on the tracks. We obtained 22 clusters when networks were clustered according to their quantitative parameter values; the value of parameters was considered in addition to their sign ***Spiess et al. (2022)***. In order to select a network for further study, we quantified signalling and T-box expression dynamics in the *in vitro* experiments, used the signalling dynamics to simulate all 22 representative networks and selected networks that also recapitulated the T-box expression dynamics measured *in vitro*. To characterise the levels and dynamics of Wnt and FGF signals *in vitro* we quantified Wnt signalling activity by performing HCR against gfp mRNA produced from a Tg(7xTCF-Xla.Siam:GFP) zebrafish reporter line which reports on active Wnt signaling ***Moro et al. (2012)*** while FGF was assayed using antibody staining against diphosphorylated ERK (Figure 4A-B). Quantification of these signals in multiple cells *in vitro* revealed that the levels of Wnt and FGF signaling decline in single cells over the 6 hours of culture. The cell population maintains high ppERK phosphorylation during the first 4 hours before downregulating it at 6 hours (Figure 4C). Mean TCF reporter levels also decline rapidly within the first two hours of culture and remain low for the remaining duration of the timecourse (Figure 4D). This was further confirmed by bulk RNA extraction and qPCR using primers for axin2 and sprouty4, which function as downstream readouts of Wnt and FGF signaling respectively (Figure 4E,F).

**Figure 4.**
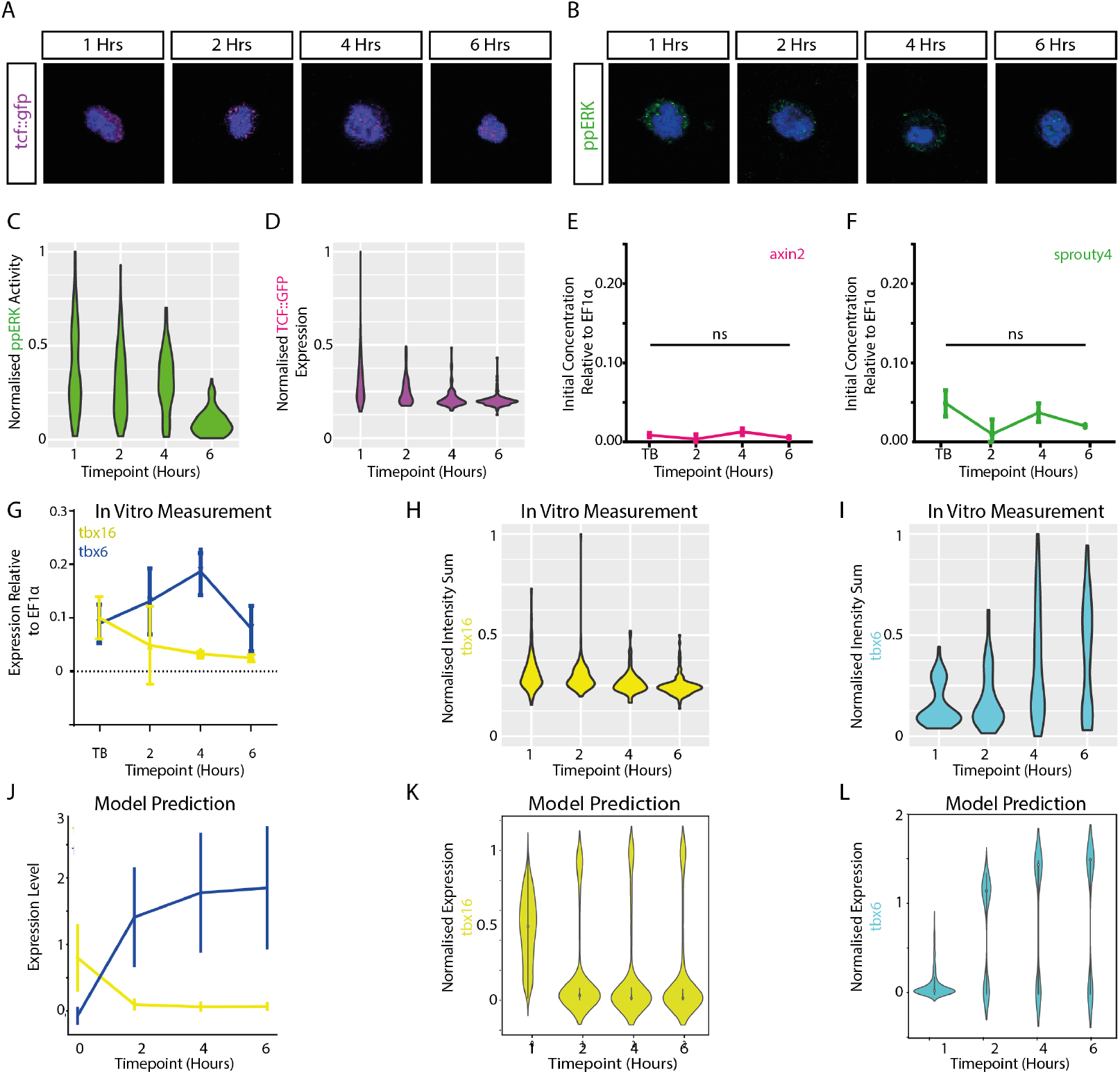
*in vitro* cultures of PSM cells differentiate as single cells with dynamics predicted by a simulated gene regulatory network. Posterior PSM cells cultured *in vitro* **(A-B)** maintain some level of Wnt and FGF signalling measured by gfp mRNA production from a 7XTCF::GFP reporter or diphosphorylated ERK antibody staining respectively. Examining these stains at the single cell level demonstrate that **(C-D)** the level of FGF and Wnt signalling declines over the time course with a gradual decline in mean ppERK activity and a more rapid decline of TCF::GFP activity over time. This is further demonstrated **(E-F)** by bulk qPCR of cultured cells over a timecourse using primers for axin2 and sprouty4 to measure levels of Wnt and FGF signalling. In the same qPCR experiments, **(G)** the level of tbx16 declines over time whilst the level of tbx6 increases to a maximum at 4 hours before then declining again. **(H-I)** Examining single cell HCRs for tbx16 and tbx6 demonstrate that the level of tbx16 declines and the level of tbx6 increases in only some cells to form a bimodal population by 6 hours, where some cells are tbx6 positive and others and still low in tbx6 expression. Using the remaining 22 reverse engineered networks which could generate patterns similar to those observed in vivo when simulated on cell tracking data, **(J)** the bulk qPCR experiment was simulated with the predicted outcome closely resembling the experimental outcome *in vivo*. **(K-L)** Simulation of single cells cultured *in vitro* and imaged as individuals also predicted the formation of a bimodal population, as was observed *in vivo*

Using bulk qPCR of cells experimentally dissociated and cultured *in vitro*, we measured a bulk loss of tbx16 mRNA and a gain of tbx6 expression over time (Figure 4G), finding tbx6 levels reaching peak expression after four hours before becoming downregulated. Next, we imaged at the single cell level *in vitro* using HCR stains against tbx16 and tbx6 experiments and quantified expression levels by masking around individual cells normalised by cell area. From this we found bimodal differentiation dynamics in single cells, where a population of cells increased their levels of tbx6 and downregulated tbx16, while at the same time, other cells remain tbx6 negative (Figure 4H,I).

Out of 22 networks, ten recapitulated both the coordinated bulk upregulation of tbx6 (Figure 4J) and its bimodal activation in single cells (Figure 4K-L). Unable to discriminate between these further based on fit, we selected a network whose predicted interactions were also supported by the current literature, namely, the network where tbx16 is predicted to activate tbx6 ***Bouldin et al. (2015)***; ***Fior et al. (2012)*** and FGF is predicted to activate tbx16 ***Goto et al. (2017)***; ***Griffin et al. (1998)*** (Figure 3C). In imposing this selection criteria, the remaining network proposed also infers that Wnt signalling will repress the expression of tbx6; an interactions which is also supported by experimental evidence ***Fior et al. (2012)***. The experimental evidence that Wnt activates expression of tbxta has also been incorporated within this model as a non-inferred prior parameter ***Bouldin et al. (2015)***; ***Goto et al. (2017)***. Taken together, these results propose a GRN (Figure 3F) that has a number of inferred and prior known interactions supported by the literature and capable of predicting differentiation dynamics of tailbud progenitors *in vivo*, on cell tracks, and when cultured as single cells *in vitro* (Figure 3E-F) and (Figure 4J,K,L).

### Cell rearrangements tune the dynamics of mesoderm progenitor differentiation *in vivo* to generate pattern emergence as a function of temporal Wnt and FGF exposure

Our live-modelling framework makes it possible to explore the differentiation dynamics of single cells within the PSM and to associate these dynamics with the cells’ relative positions and movements, linking them back to the Wnt and FGF signalling environments that they each experience as morphognesis unfolds. In particular, it allows us to explore how cell movements might be playing a role controlling the timing of tbx6 onset as a function of Wnt and FGF exposure by determining when the cells exit from the progenitor domain. This would in turn help explain the range of differentiation timescales observed *in vivo* (Figure 2A-F, M).

We investigate this further by choosing two progenitor cells that both begin the simulation at 23% posterior-anterior position within the progenitor region, but that as development proceeds, will diverge notably in their positions and overall 4D trajectories: one cell will leave the progenitor region and enter the PSM, making its way towards a somite Figure 5A, while the other will remain in the progenitor region even moving slightly posteriorly over time (Figure 5D). As a result of these different 3D movements over time, the cells’ signalling AGETS are different Figure 5B,E. The first thing to note is that the signalling AGETs are initially not the same for both cells despite them being located at the same anteroposterior position. This reflects the fact that AGETs are constructed from 3D gene expression quantification and retain the heterogeneity present in the data, which is present even across narrow spatial domains ***Spiess et al. (2022)***. The next thing to note, is that the cell that exits the progenitor region will continue to upregulate FGF and downregulate Wnt (Figure 5B), while the cell that remains in the progenitor region will maintain high Wnt signalling levels and low FGF (Figure 5E).

**Figure 5.**
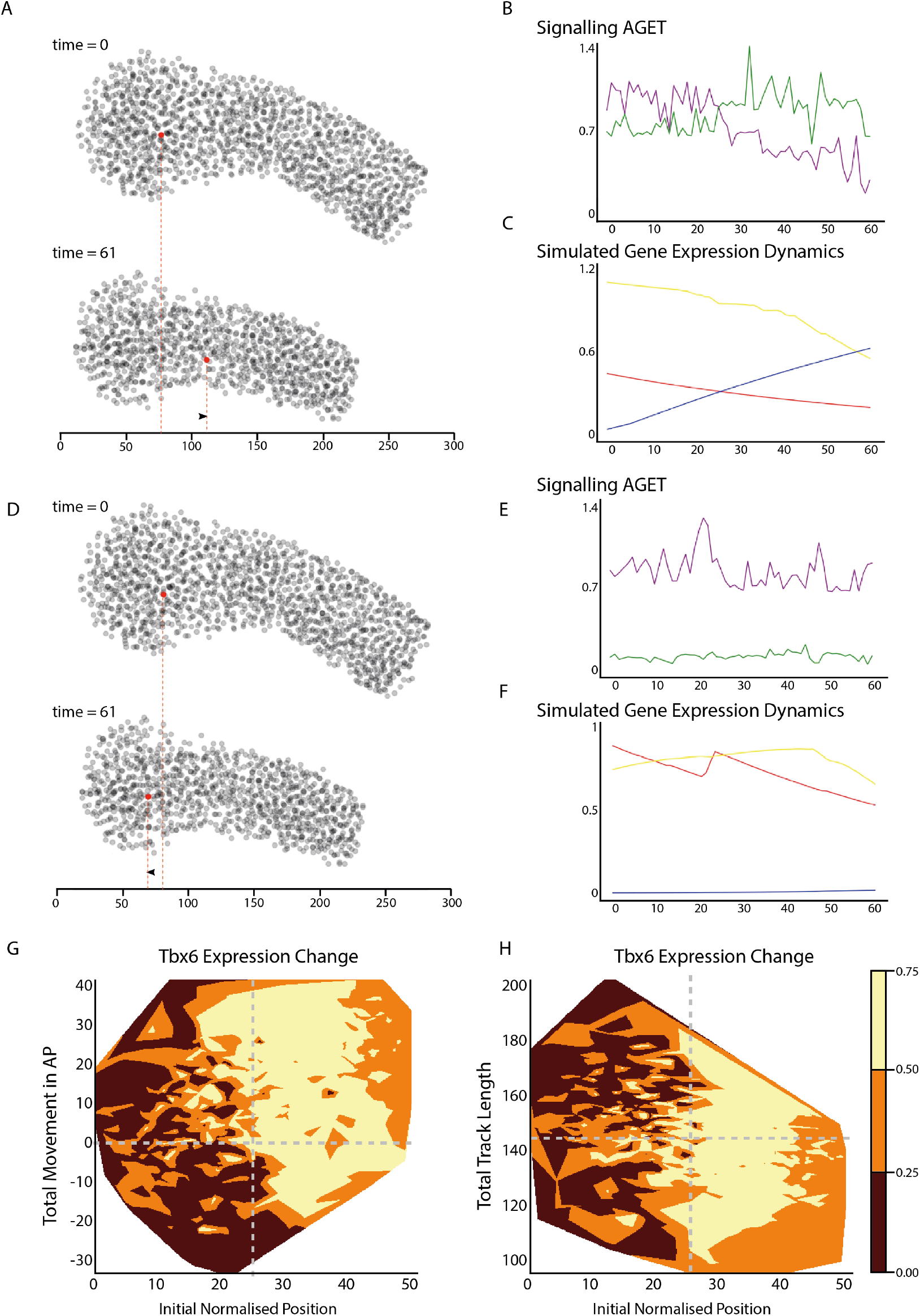
Temporal signalling dynamics implemented by cell movements and rearrangements control the timing of differentiation in mesoderm progenitors. **(A. top)** Relative position of the cell within the zebrafish PSM at the start of the simulation (23.8321% posterior-anterior at time = 0) and **(A. bottom)** at the end of the simulation (time = 61). **(B)** Signalling AGET and **(C)** simulated T-box gene expression dynamics for the cell shown in (A). **(D. top)** Relative position of the cell within the zebrafish PSM at the start of the simulation (23.8361% posterior-anterior at time = 0) and **(D. bottom)** at the end of the simulation (time = 61). **(E)** Signalling AGET and **(F)** simulated T-box gene expression dynamics for the cell shown in (D). **(G)** Visualisation of the degree of change in Tbx6 gene expression over the simulation compared to total cell movement in the direction of the anterior-posterior axis for cells starting in the posterior 25% region of the PSM. Positive Total Movement in AP represents anterior cell movement whereas negative Total Movement in AP represents posterior cell movement. Cells have the largest change in tbx6 expression when displacing anteriorly. Aberrant tbx6 expression in posterior moving cells can be observed in the bottom left quadrant where cells are moving towards the posterior yet upregulating tbx6 expression, shown by the light ochre colouring. **(H)** Visualisation of the degree of change in Tbx6 gene expression over the simulation compared to total track length for cells starting in the posterior 25% region of the PSM.The total distance a cell travels shows no correlation with the degree of tbx6 gene expression change as top and bottom quadrants show similar degrees of ochre shading.

As a result of their different patterns of movement, and therefore different Wnt and FGF dynamics, the simulated T-box gene expression dynamics differ in both cells. The cell which exited the progenitor region is predicted by the model to reduce its tbxta and tbx16 expression levels and slowly increase tbx6 expression (Figure 5A,C). By contrast, the cell which remains embedded in the progenitor region throughout the simulation, is predicted to maintain high levels of tbxta and tbx16 expression, and low level of tbx6 expression (Figure 5D,F). In either case, the associated Wnt and FGF profiles are modified due to the different rate of movement through the 3D tissue level signalling environments, also taking into account their heterogeneity within the tissues.

In addition our model suggests that it is a progenitor cell’s overall movement along the anterior-posterior axis rather than the total distance it travels (track length) that determines its differentiation dynamics. Figure 5G shows that cells starting in the posterior-most 25% region of the progenitor region will experience a larger change in tbx6 expression levels if their overall displacement is positive (towards the anterior) than negative (towards the posterior), as reflected by the larger lightly-coloured area in the top left quadrant as compared to the bottom left one. In contrast, the total distance travelled by a cell in the same region does not predict or correlate with the level of tbx6 change that it will experience, which is shown by the comparable shading in both top and bottom left quadrants in Figure 5H.

All in all we show that in our model cell movements themselves are responsible for modulating the dynamics of progenitor differentiation by generating dynamic exposure to spatial signalling environments, and expression patterns emerge as a result at the tissue level through a combination of dynamic signalling exposure, cell intrinsic gene regulatory interactions, and cell movements.

### Posterior cell displacements predict increased tbx6 heterogeneity within the tail-bud

Figure 5G and H also reveal a degree of heterogeneity in the level of tbx6 expression change within cells starting in the posterior-most 25% of the embryonic tailbud. This is reflected by the existence of patches of all three tones of orange in the top and bottom left quadrants of Figure 5G and H. In particular, the presence of light ochre patches in the bottom left quadrant of Figure 5G suggests that a subset of cells upregulate tbx6 without leaving the progenitor domain.

As a result, when simulated on live tracking data, our network predicts a degree of tbx6 expression heterogeneity within the tailbud (Figure 6A-B). To identify whether there is indeed a posterior bias of ectopic expression of *tbx24* as predicted by the live modelling, the nuclei within the PSM were segmented (Figure 6C) and then classified as either anterior or posterior relative to the posterior end of the notochord within each embryo (Figure 6D). Following this, cells with aberrant gene expression were identified in each domain, defined as expressing *tbx6* in the posterior, or *tbx16* in the anterior domain (Figure 6E). The frequency of aberrant gene expression was measured and we confirmed a posterior bias in erroneous gene expression, as predicted by the live modelling (Figure 6F) indicating that although this system is able to generate deterministic patterns at the level of the tissue (Figure 2D-F), it does so from noisy gene expression at the single cell level, as seen *in vivo* (Figure 6G).

**Figure 6.**
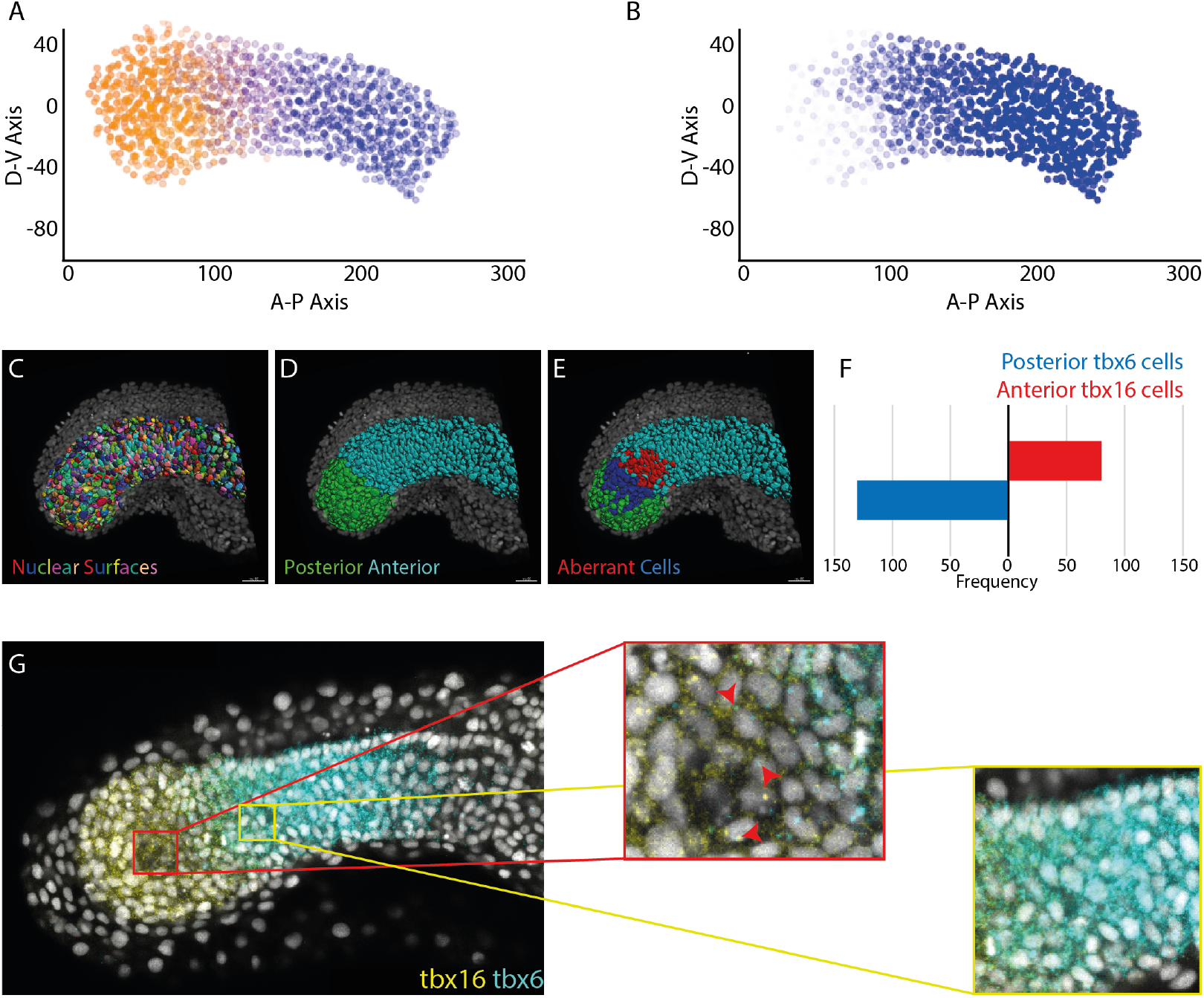
Live modelling predicts cell level heterogeneity. **(A-B)** Live modelling simulations predict that some cells upregulate tbx6 in the posterior domain of the PSM, where tbx16 would be expected to be expressed by the end of the simulation. (A) All three genes are shown: tbxta in red, tbx16 in yellow and tbx6 in blue. (B) Same as (A) showing only tbx6 expression in blue. **(C-E)** By examining whole mount HCR images and segmenting individual nuclei, cells could be classified as either posterior or anterior relative to the end of the notochord. **(F)** Cells in each of these domains could then be classified and the number of cells expressing tbx6 in the tbx16 domain and vice versa counted. This demonstrates a posterior bias in aberrant gene expression as predicted by the live modelling simulations. **(G)** These aberrant cells can be identified in slices of HCR data.

## Conclusions

During development, cells undergo state transitions at a rate that is in part determined by cell intrinsic timers of differentiation. Mechanistically, this developmental tempo is set by the time it takes for the underlying GRN to elicit changes in gene expression. It is to a large degree also dependent on protein production and degradation rates within each cell ***Matsuda et al. (2020)***; ***Rayon et al. (2020)***, and species can vary these rates in a manner that is linked to their basal metabolic rate ***Diaz-Cuadros et al. (2021)***. In a multi-cellular context, extrinsic signals can further tune the dynamics of this process by controlling the activation or inhibition of other nodes within a GRN, leading to the creation of a non-autonomous dynamical system where the state change of a cell is directly linked to the strengths of signal inputs that it encounters ***Busby and Steventon (2020)***. Dynamical systems models appropriately account for both intrinsic and extrinsic timers of differentiation and are therefore a powerful tool to probe how gene expression patterns emerge during development ***Crombach et al. (2016)***; ***Verd et al. (2014)***; ***Sagner and Briscoe (2017)***; ***Kicheva et al. (2014)***. Here, we present a GRN that can explain the temporal dynamics of PSM progenitor differentiation *in vitro*, and crucially is also able to recapitulate the emergence of tissue-level T-box pattern formation *in vivo* in the context of cell rearrangement.

Probing the predicted dynamics of gene expression change for single cells within our model has revealed two insights. Firstly, an unexpected degree of heterogeneity observed for tbx6 expression in the posterior progenitor domain. Gene expression heterogeneity is a common feature of progenitor populations that has in many cases been linked to underlying stochasticity in gene expression at the transcriptional level ***Moris et al. (2016)***. While this is also likely to be the case for T-box genes in the PSM, our model points to a second source of noise in the system. GRNs are configured in such a way that produces a temporal delay in response to morphogen exposure, as such cells will not immediately adjust their expression state in situations where they move into an altered signalling environment. For the PSM, this means that cells displaced posteriorly from a tbx6 positive region into the tbx16 progenitor domain retain their previous gene expression state and thereby contribute to gene expression heterogeneity. Dissecting these multiple sources of heterogeneity in developing systems will be an important step in fully exploring the role of gene expression noise *in vivo*.

A second insight derived from our model relates to the role of cell movements in the emergence of tissue-wide gene expression patterns. To generate patterns of T-box expression in the PSM, cells must tune their intrinsic dynamics of differentiation to the rate at which they exit the tailbud and enter the PSM. We observe that cells delayed in their exit from a high tbxta expressing progenitor state encounter higher Wnt levels and do not transit through to the lower region of FGF activity in the PSM. As such, cell movements determine the level and dynamics of morphogen exposure, and in doing so regulate their differentiation rate accordingly. This demonstrates a crucial role for cell movements in the emergence of gene expression pattern that is a composite of morphogen exposure, intrinsic timers of differentiation, and cell rearrangement. It emphasises the difficulty in determining gene function from mutant phenotype analysis alone as a given signal or transcription factor may have dual impact on gene expression patterns, firstly through the regulation of other network nodes to control their levels of expression but also indirectly via the regulation of cell movements within the tissue in question. This is especially important in the context of PSM development where signals such as Wnt and FGF have known functions in also regulating epithelial-to-mesenchymal transitions ***Bajard et al. (2014)***; ***Goto et al. (2017)***; ***Row et al. (2011)***. In addition, tbx16 has a clear role in both the specification of mesodermal cell fate and the control of cell movement from the tailbud ***Griffin et al. (1998)***. Here, we present a novel approach to model this highly dimensional system, that considers both GRN interactions and cell movements. We envisage that this will be a powerful predictive tool when testing proposed network interactions in the context of cell tracking data obtained from both wild-type and mutant embryos.

Embryonic development is characterised by a series of multi-scalar interactions where dynamic state changes of individual cells, when coupled with cell movements and morphogen exposure, can lead to the emergence of gene expression patterns at the tissue level. Cell movements themselves are likely to also be impacted by tissue level properties such as the liquid-to-solid transition observed in the PSM ***Mongera et al. (2018)***; ***Serwane et al. (2017)***, and furthermore via forces acting between adjacent tissues during the process of posterior body elongation ***McLaren and Steventon (2021)***; ***Thomson et al. (2021)***; ***Steventon et al. (2016a)***; ***Tlili et al. (2019)***. As such, it is essential that we approach the role of GRNs in development in a context of multi-tissue morphogenesis to gain a complete picture development. The live-modelling framework utilised here provides an inroad into achieving this as it enables predictions to be generated of how specific GRN topologies can lead to pattern emergence in multiple morphogenetic contexts. This might be in the context of interpreting complex mutant phenotypes or to probe how developmental systems respond to perturbations at the level of physical tissue properties, tissue geometry or multi-tissue mechanical interactions. Furthermore, while in many cases it is assumed that evolution acts to drive changes in gene expression through altering GRN interactions, our work points to an equivalent potential in altering cell movements within a tissue in question while conserving the GRNs. Exploring the multi-dimensional regulation of evolutionary change is therefore an important direction for the field.

## Materials and Methods

### Animal Husbandry

This research was regulated under the Animals (Scientific Procedures) Act 1986 Amendment Regulations 2012 following ethical review by the University of Cambridge Animal Welfare and Ethical Review Body (AWERB). Embryos were obtained and raised in standard E3 media at 28°C. Wild Type lines are either Tüpfel Long Fin (TL), AB or AB/TL. The Tg(7xTCF-Xla.Sia:GFP) reporter line ***Moro et al. (2012)*** was provided by the Steven Wilson laboratory. Embryos were staged as in ***Kimmel et al. (1995)***

### Primary Culture of Tailbud Progenitor Cells

Cells were explanted from the tailbud as in ***Webb et al. (2016)***; ***Rohde et al. (2021)***. Effort was made to remove the ectoderm prior to dissection. Cells were dissected in calcium and magnesium free PBS in order to promote cell dissociation. Cells were cultures in 8 well Ibidi Micro-Slides under the fully defined L15 media supplemented with PenStrep solution to limit bacterial growth.

RNA extractions were made in triplicate, from independent experiments, using Trizol Reagent (Ambion LifeTechnologies) following a standard protocol and reverse transcription using Superscript III (Invitrogen). Resultant cDNA was quantified using SYBRGreen with liquid handling robot (Qiagility, Qiagen) and analysed on a RotorGeneQ thermocycler (Qiagen).

Primer sequences:

*axin2* 5’-TACCCTCGGACACTTCAAGG-3’ and 5’-TGCCCTCATACATTGGCAGA-3’; *sprouty4* 5’-CACGCGCCCTAGTATCAAAC-3’ and 5’-GGGATCTTGGTGAAGTGTGC-3’; *EF1a* 5’-GGAGACTGGTGTCCTCAA-3’ and 5’-GGTGCATCTCAACAGACTT-3’.

Concentration of cDNA was estimated using an in-house MAK2 analysis method, as described previously ***Turner et al. (2014)***.

### In Situ Hybridisation Chain Reaction (HCR)

Embryos were raised to the required stage then fixed in 4% PFA in DEPC treated PBS without calcium and magnesium at 4°C overnight. Embryos were then stained using HCR following the standard zebrafish protocol found in ***Choi et al. (2018)***. Probes, fluorescent hairpins and buffers were all purchased from Molecular Instruments. After staining, samples were counter stained with DAPI and mounted under 80% glycerol.

### Immunohistochemistry

Embryos were raised to the required stage then fixed in 4% PFA in DEPC treated PBS without calcium and magnesium at 4°C overnight. Embryos were then blocked in 3% goat serum in 0.25% Triton, 1% DMSO, in PBS for one hour at room temperature. Diphosphorylated ERK was detected using the primary antibody (M9692-200UL, Sigma) diluted 1 in 500 in 3% goat serum in 0.25% Triton, 1% DMSO, in PBS. The samples were incubated at 4°C overnight then washed in 0.25% Triton, 1% DMSO, in PBS. Secondary Alexa 647nm conjugated antibodies were diluted 1 in 500 in 3% goat serum in 0.25% Triton, 1% DMSO, 1X DAPI in PBS and applied overnight at 4°C.

### Imaging and Image Analysis

Samples were imaged using a Zeiss LSM700 inverted confocal microscope at 12 bit, 20X or 40X magnification, with an image resolution of 512×512 pixels. Single cell HCR was imaged using a Nikon Ti inverted widefield microscope at 63X magnification.

Image analysis of confocal images was done using the line drawing tool on Fiji ***Schindelin et al. (2012)***; ***Schneider et al. (2012)*** set to a width of 50 pixels. Lines were drawn following the curve of the embryo, through the centre of the PSM from posterior PSM to the posterior most clear somite boundary. Profiles were normalised to the length of the PSM and signal intensity as individual embryos by dividing the measured value by the maximum value of that embryo.

Nuclear segmentation of whole embryos stained using HCR was conducted using a tight mask applied around the DAPI stain using Imaris (Bitplane) with a surface detail of 0.5µm. Touching surfaces were split using a seed size of 4µm. Values were exported as X, Y, Z coordinates relative to the posteriormost tip of the PSM where X, Y, Z were equal to (0, 0, 0). The PSM was then segmented by hand by deleting nuclear surfaces outside of the PSM, including notochord, spinal cord, anterior somites and ectoderm. Only the PSM closest to the imaging objective, therefore of highest imaging quality was measured with the distal PSM also removed.

Intensity mean values of each transcription factor HCR signal were exported and normalised between 0 and 1 by dividing each cell’s mean signal intensity by the maximum measured within that sample, per gene. PSM length was normalised individually between 0 and 1 by division of the position in X by the maximum X value measured in each embryo.

Single cell image analysis was conducted using Imaris (Bitplane) by generating loose surface masks around the DAPI stain to capture the full nuclear region and cytoplasm. Surface masks were then filtered to remove any masks where two cells joined together or small surfaces caused by background noise, or fragmented apoptotic nuclei. The intensity sum of each channel was measured and normalised by the area of the surface, as surface area and transcript intensity had been demonstrated to correlate. Expression level was then normalised between 0 and 1 using the maximum value measured for each gene, in each experiment.

Live imaging datasets of the developing PSM was created using a TriM Scope II Upright 2-photon scanning fluorescence microscope equipped Insight DeepSee dual-line laser (tunable 710-1300 nm fixed 1040 nm line). Embryo was imaged with a 25X 1.05 NA water dipping objective. Time step and frame number as per figure legend. Embryos laterally in low melting agarose with the entire tail cut free to permit normal development ***Hirsinger and Steventon (2017)***.

### Model formulation

We formulated the T-box gene regulatory network using a dynamical systems formulation. The models aim is to recapitulate the dynamics of T-box gene expression for any cell, or rather a general cell, in the developing zebrafish PSM. We use a connectionist model formulation previosly used to model other developmental patterning processes ***Mjolsness et al. (1991)***.

The mRNA concentrations encoded by the T-box genes *tbxta, tbx16* and *tbx24* are represented by the state variables of the dynamical system. For each gene, the concentration of its associated mRNA *a* at time *t* is given by *g*^*a*^(*t*). mRNA concentration over time is governed by the following system of three coupled ordinary differential equations:

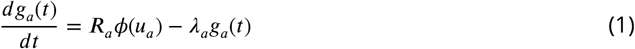

where *R*^*a*^ and *λ*^*a*^ respectively represent the rates of mRNA production and decay. *ϕ* is a sigmoid regulation-expression function used to represent the cooperative, saturating, coarse-grained kinetics of transcriptional regulation and introduces non-linearities into the model that enable it to exhibit complex behaviours:

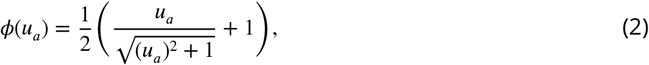

where

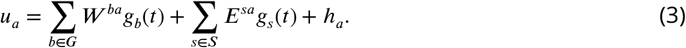

*G* = {*tbxta, tbx*16, *tbx*24} is the set of T-box genes while *S* = {Wnt, FGF} is the set of external regulatory inputs provided by the Wnt and FGF signalling environments. The concentrations of the external regulators *g*_*s*_ are interpolated from quantified spatial mRNA expression data (Figure 1J) and translated into time as explained in the main text to used as dynamic inputs to the model. Changing Wnt and FGF concentrations over time renders the parameter term 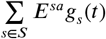 time-dependent and therefore the model non-autonomous ***Collier et al. (1996)***; ***Verd et al. (2014)***.

The interconnectivity matrices *W* and *E* house the parameters representing the regulatory interactions among the T-box genes, and from Wnt and FGF to the T-box genes, respectively. Matrix elements *w*^*ba*^ and *e*^*sa*^ are the parameters representing the effect of regulator *b* or *s* on target gene *a*. These can be positive (representing an activation from *b* or *s* onto *a*), negative (repression), or close to zero (no interaction). *h*_*a*_ is a threshold parameter representing the basal activity of gene *a*, which acknowledges the presence of regulators absent from our model. Model parameters are detailed in Table S1.

### Model fitting and selection

We have developed a methodology that makes it possible to reverse-engineer gene regulatory networks that might be driving 4D tissue-level pattern formation during morphogenesis ***Spiess et al. (2022)***. Prior to the development of this methodology, it was only possible to confidently reverse-engineer GRNs underlying pattern formation in developmental processes where the timescales of pattern formation and morphogenesis could be separated. This was because it is only possibly to accurately quantify the gene expression dynamics of multiple genes of interest in a single cell (or at a given position in the tissue) from static confocal stains of the tissue at different developmental stages if the cells are not significantly changing their position over time. Otherwise, the quantification at a given position in the tissue reflects the gene expression dynamics of many cells, and not that of a single cell, which is what the GRN is modelling, leading to inaccurate GRN predictions. This limitation has hence historically greatly restricted the types of patterning processes that could be reverse-engineered.

In order to extend the application of reverse-engineering approaches to a wider variety of patterning processes -particularly those where the timescales of pattern formation and morphogenesis are coupled - we have developed a novel methodology that is based on approximating gene expression trajectories of single cell tracks and using these for gene regulatory network inference ***Spiess et al. (2022)***. In brief, this methodology can be split into two main parts: generating approximated gene expression trajectories (AGETs) for every cell track, and using a Markov Chain Monte Carlo parameter sampling algorithm to infer candidate GRNs that can recapitulate the gene expression dynamics of a subset of the AGETs. Candidate GRNs are then selected for further study based on their ability to recapitulate the tissue-level patterning dynamics, and other case-specific factors ***Spiess et al. (2022)***.

In order to construct the AGETs we align and project gene and signalling quantifications obtained from confocal imaging HCR and immuno stained tailbuds onto each time frame in the timelapse of the developing PSM. Each cell in the time frame is assigned gene expression and signalling levels averaged from the five closest cells to it from the quantification. This is repeated for each frame, and results in an approximated gene and signalling trajectory for each cell in the movie. Ten AGETs were then selected pseudo-randomly along the PSM and used for network inference. The MCMC algorithm was run 100 times, and solutions were resolved into 22 clusters that were able to recapitulate tissue-level T-box patterning when simulated on the tracks. Representative networks of these 22 clusters were then used to simulate the dissociation experiments and one network was selected for further study. For further details of this methodology please refer to the methodology paper ***Spiess et al. (2022)***.

## Acknowledgments

We would like to thank Buzz Baum, Laurel Rohde, Andrew Oates and the team, Dillan Saunders and Meagan Hennessy for feedback on this manuscript. We would like to thank the Steven Wilson lab for sharing the Tg(7xTCF-Xla.Sia:GFP) line, and Kawamura Akinori for the Tbx6::GFP line. Thanks to the Cambridge Advanced Imaging Centre (CAIC) for imaging support. T.F., S.H. and B.S. are supported by a Henry Dale Fellowship jointly funded by the Wellcome Trust and the Royal Society (109408/Z/15/Z) and T.F. by a scholarship from the Cambridge Trust, University of Cambridge. L.T. is supported by a scholarship from the BBSRC. Y.W. is supported by a summer vacation stipend from St Catharine’s College, University of Cambridge. B.C. is supported by a Stipend from the Bedford Fund, King’s College, University of Cambridge and a scholarship from the Wellcome Trust. B.V. was supported by a Herschel Smith Postdoctoral Fellowship, University of Cambridge and Department of Zoology, University of Oxford. B. C. is supported by a Wellcome Trust Developmental Mechanisms PhD studentship (222279/Z/20/Z). The work was supported by Wave 1 of the UKRI Strategic Priorites Fund under the EPSRC grant EP/T001569/1, particularly the “AI for Science and Government” theme within that grant and the Alan Turing Institute.

## Notes

### Competing Interest Statement

The authors have declared no competing interest.

